# A flexible model for thermal performance curves

**DOI:** 10.1101/2024.08.01.605695

**Authors:** Mauricio Cruz-Loya, Erin A Mordecai, Van M Savage

## Abstract

Temperature responses of many biological traits—including population growth, survival, and development—are described by thermal performance curves (TPCs) with phenomenological models like the Briere function or mechanistic models related to chemical kinetics. Existing TPC models are either simple but inflexible in shape, or flexible yet difficult to interpret in biological terms. Here we present flexTPC: a model that is parameterized exclusively in terms of biologically interpretable quantities, including the thermal minimum, optimum, and maximum, and the maximum trait value. FlexTPC can describe unimodal temperature responses of any skewness and thermal breadth, enabling direct comparisons across populations, traits, or taxa with a single model. We apply flexTPC to various microbial and entomological datasets, compare results with the Briere model, and find that flexTPC often has better predictive performance. The interpretability of flexTPC makes it ideal for modeling how thermal responses change with ecological stressors or evolve over time.

## Introduction

A fundamental problem in ecology is to understand how the growth, physiology, and behavior of organisms depend on their environment. Temperature variation is an important environmental characteristic due to its multiple effects on the physiology (Knapp & Huang 2022) and behavior (Ito & Awasaki 2022) of organisms. Through these effects, changes in temperature impact the fitness of organisms (Amarasekare & Savage 2012) and ultimately the distribution of species across geographic space (Jeffree & Jeffree 1994). Understanding the effects of temperature in organisms is thus crucial to predict how climate change will modify the geographic distribution of species and their interactions, as well as its potential impacts on biodiversity (Nunez *et al*. 2019; Waldock *et al*. 2018), agriculture (Jägermeyr *et al*. 2021), the transmission of infectious disease (Rocklöv & Dubrow 2020), and other important ecosystem processes.

Many traits, including rates of metabolism (Schulte 2015), population growth (Savage *et al*. 2004), and development (Briere *et al*. 1999) vary continuously and nonlinearly with temperature. This dependence can be represented by a thermal performance curve (TPC) that describes the value or performance of the trait at different temperatures (Huey & Kingsolver 1989). Empirically, TPCs are often unimodal, reaching maximum performance at a single optimum temperature and decreasing to a thermal maximum and minimum where performance goes to zero (Angilletta Jr. 2009; Dell *et al*. 2011; Huey & Berrigan 2001).

Various mathematical models have been developed to describe TPCs quantitatively (Arroyo *et al*. 2022; Briere *et al*. 1999; Hultin *et al*. 1955; Johnson & Lewin 1946; Ratkowsky *et al*. 1983, 2005; Ritchie 2018; Schoolfield *et al*. 1981; Sharpe & DeMichele 1977; Shi & Ge 2010; Yin *et al*. 1995). These models make it possible to infer useful summaries of the temperature dependence of a trait (such as the optimum, maximum, and minimum temperatures) from experimental data. These summaries can then be compared between different populations of the same species, across species, or across traits (Barton & Yvon-Durocher 2019; Bennett 1980; Buckley & Huey 2016; Couper *et al*. 2024; Gounot 1976; Knies *et al*. 2009; Shocket *et al*. 2020). Models of TPCs are also used as building blocks in more complex mathematical models that describe population dynamics and interactions between species. For instance, due to the sensitivity of ectotherm physiology to environmental temperature, transmission dynamics of vector-borne diseases are often highly sensitive to temperature. Mathematical models for the temperature-dependent transmission of these diseases can be constructed using TPC models for traits of the vector, host, and pathogen that affect disease transmission (Mordecai *et al*. 2013, 2017; Shocket *et al*. 2020). Models of predator-prey dynamics that incorporate the effects of temperature are also based on TPC models for traits of the prey and predator (Dell *et al*. 2014; Gilbert *et al*. 2014; Pepi *et al*. 2023).

Thermal performance models can broadly be classified into mechanistic models that derive from an underlying theory (Arroyo *et al*. 2022; Hultin *et al*. 1955; Johnson & Lewin 1946; Ratkowsky *et al*. 2005; Ritchie 2018; Schoolfield *et al*. 1981; Sharpe & DeMichele 1977) and phenomenological models that fit empirical data without attempting to explain the underlying mechanism that gives rise to the TPC (Briere *et al*. 1999; Logan *et al*. 1976; Ratkowsky *et al*. 1983; Yin *et al*. 1995). Mechanistic models have some advantages, as they can be used to link TPCs to other biological traits, such as body size or metabolic rate through theoretical frameworks like the metabolic theory of ecology (Kirk *et al*. 2018; Molnár *et al*. 2013, 2017; Savage *et al*. 2004). However, mechanistic TPC models are often parametrized in terms of quantities that can be difficult to interpret in ecological terms (e.g., the activation energy for a potentially rate-limiting chemical reaction for the trait being measured). Because of this, many ecological and epidemiological applications use phenomenological models that are parametrized in terms of more interpretable quantities (such as maximum and minimum temperatures) while still providing a good fit to experimental data, often with fewer parameters than mechanistic models. Moreover, many phenomenological models have explicit thermal limits for trait performance rather than an asymptotic decrease, which is desirable for modeling some traits (e.g., probability of survival to adulthood).

One popular set of phenomenological models—the Briere models—are commonly used to describe the temperature dependence of insect developmental rates (Briere *et al*. 1999) and have been widely adopted in the ectotherm thermal biology literature (Haye *et al*. 2014; Lachenicht *et al*. 2010; Lemoine 2017; Mordecai *et al*. 2013, 2017; Paaijmans *et al*. 2009; Sentis *et al*. 2012; Tochen *et al*. 2014). These models are based on the same mathematical equation (Equation 1), differing only in the number of free parameters. The sparser three-parameter model—commonly referred to as the Briere1 model (or just the Briere model)—is popular in applications due to its parsimony, the biological interpretability of two of its parameters (the minimum and maximum temperatures), and its ability to describe many left-skewed TPCs for biological rates (Briere *et al*. 1999; Mordecai *et al*. 2013, 2017).

However, both Briere models have shortcomings that should be carefully considered before their use. First, the Briere1 model makes a very strong implicit assumption about the relationship between the minimum, maximum, and optimum temperatures that does not have a biological justification and that can potentially bias optimum temperature estimates. Second, due to their mathematical structure, the Briere1 and Briere2 models cannot describe thermal performance curves from psychrophilic organisms that can function below freezing temperatures. Lastly, the Briere models can only describe thermal performance curves that are left-skewed but are unable to describe TPCs with different shapes. This limitation is important when the goal is to compare traits that differ in TPC shape, such as symmetric and asymmetric responses.

As an alternative to the Briere models we present a flexible model for thermal performance curves that addresses these limitations, and can describe left-skewed, symmetric, and right-skewed unimodal TPCs of varying thermal breadth. This model, which we call flexTPC, is mathematically equivalent to the Beta model for crop development as originally presented by (Yin *et al*. 1995), which has not been widely adopted ectotherm animal physiology and ecology literature, but is reparametrized in terms of biologically interpretable quantities to make it more suitable for applications in ecology and infectious disease modeling. A previous version of this model was derived in (Cruz-Loya *et al*. 2021) by modifying the Briere2 model (Equation 1) with the goal of describing TPCs of bacterial growth under antibiotics. However, this previous work focused primarily on how antibiotics modify TPCs rather than on the much broader potential applications of the mathematical model, and the model as presented previously had a remaining parameter without a direct biological interpretation.

In this work, we provide a novel, fully biologically interpretable parametrization of the flexTPC model and compare its predictive performance with that of the Briere1 and Briere2 models in real-world datasets. We find that flexTPC has similar or better performance than the Briere models when describing insect development data, while performing much better when describing thermal performance curves of psychrophilic organisms and TPCs that are symmetric or right-skewed. Finally, we show that flexTPC can accurately describe many different mosquito life history traits for which different functional forms (linear, quadratic, and Briere1) were used in the past. Our results show that flexTPC is a flexible and interpretable descriptive model for unimodal TPCs that has some important advantages compared to the Briere models, and that is especially well-suited for applications where TPCs of different shapes need to be compared. Its interpretability is well-suited for Bayesian approaches for parameter inference, enabling the use of informative prior distributions based on biological knowledge such as the thermal range of the species habitat and typical maximum trait values for the same trait in related species.

## Methods

### The Briere models

Thermal performance curve models describe trait performance *r* as a function of temperature *T*. The Briere2 model is defined as follows:

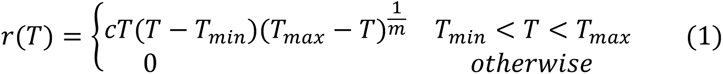

where *T_min_* and *T_max_* are the minimum and maximum temperatures for the trait, respectively, and *c*, *m* ≥ 0 are arbitrary constants. The Briere1 model is the special case of equation (1) where *m* = 2. In general, *r*(*T*) has three roots (values of *T* where *r*(*T*)=0), with one at *T* = 0°*C*. This makes the Briere models unsuitable to describe TPCs of organisms that have nonzero performance below freezing temperatures. Because of this, the Briere models are restricted to *T_min_* ≥ 0°*C* so that there are only two roots (*T_min_* and *T_max_*).

The optimum temperature of the Briere models is given by the following expression (Briere *et al*. 1999):

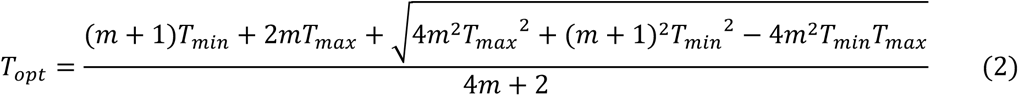

For the Briere1 model (where *m* = 2 is fixed), *T_opt_* is a deterministic function of *T_min_* and *T_max_*. In other words, it is impossible to vary *T_opt_* when *T_min_* and *T_max_* are fixed: the Briere1 model implicitly assumes a strong relationship between these parameters. To our knowledge, this assumption has no biological basis, and as a result, enforcing it will lead to biased inference of these parameters.

### The flexTPC model

The flexTPC model is defined as:

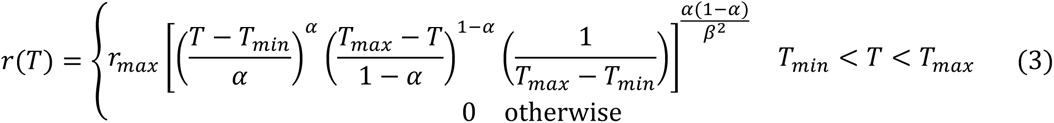

where *r_max_* is the maximum performance/value of the trait, and *T_min_* and *T_max_* are the minimum and maximum temperatures, respectively. These three parameters determine the scaling of the TPC in the temperature and performance axes (Figure 1, right panel). Two additional parameters determine the shape of the curve. Parameter *α* ∈ [0,1] determines the location of the temperature optimum *T_opt_* relative to the maximum and minimum through the relationship

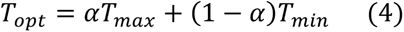

**Figure 1.**
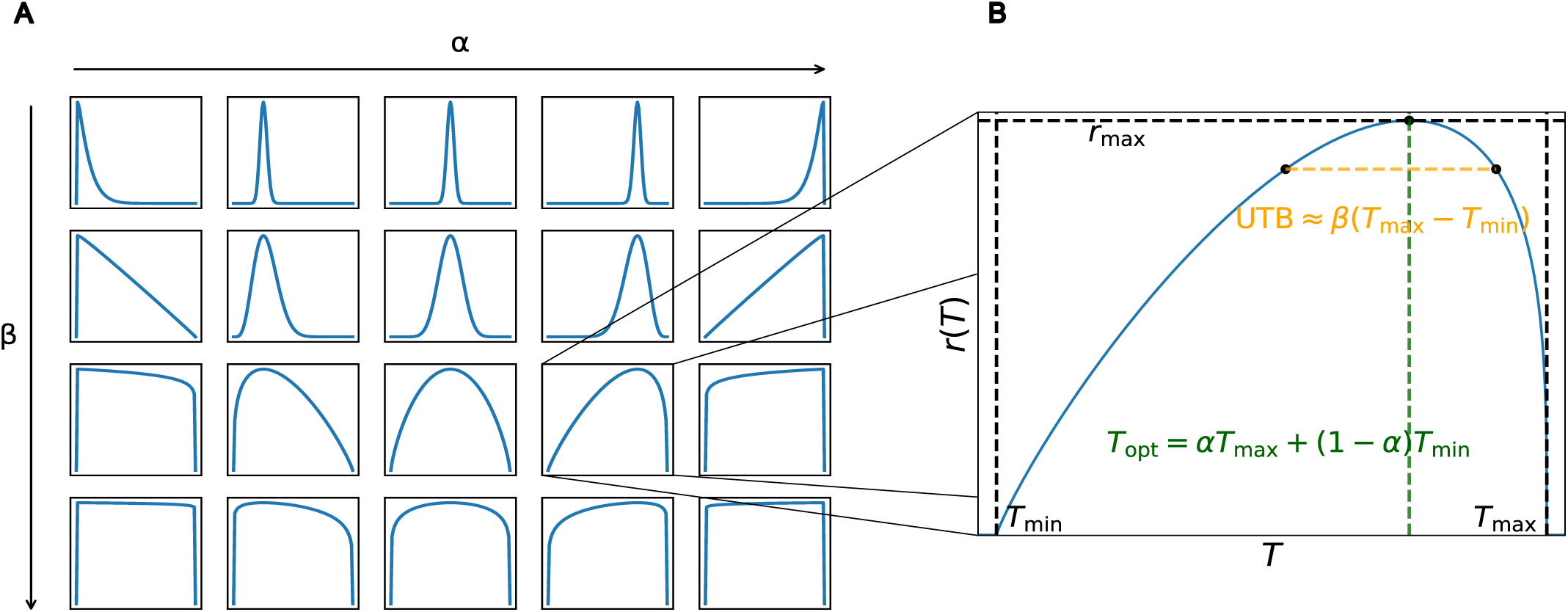
The flexTPC model can describe unimodal thermal performance curves of various shapes. *A.* The flexTPC model (Equation 3) has two parameters that determine the shape of the curve: *α* (varying from left to right) corresponds to the position of the temperature optimum relative to the minimum and maximum temperatures while *β* (varying from top to bottom) determines the thermal breadth near the top of the curve. *B.* Three additional parameters determine how the curve is scaled in the temperature and trait performance axes: the minimum and maximum temperatures (*T_min_* and *T_max_*, respectively), and the maximum value of the response *r_max_*. The optimum temperature *T_opt_* can be at any point between *T_min_* and *T_max_* : its position is determined by parameter *α* ∈ [0,1]. The upper thermal breadth (UTB), defined as the temperature range where 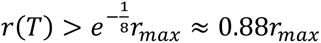, is approximately the product of *β* and the lower thermal breadth *T_max_* − *T_min_* where *r*(*T*) > 0 (for details on the accuracy of this approximation, see Methods and Figure S4).

This makes it possible for flexTPC to describe unimodal curves of any skewness by varying *α*, where e.g. *α* = 0.5 corresponds to a symmetric curve, and *α* = 0 and *α* = 1 correspond to *T_opt_* = *T_min_* and *T_opt_* = *T_max_*, respectively.

The parameter *β* > 0 determines the upper thermal breadth (UTB) of the TPC, with larger values corresponding to broader curves and smaller values to narrower curves. UTB, defined here as the temperature range for which 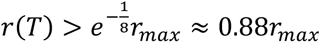 (see Supplemental Information), is approximately

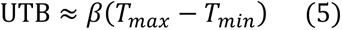

As *T_max_* − *T_min_* corresponds to the thermal breadth of nonzero performance (defined here as the lower thermal breadth), *β* is the (approximate) ratio of the upper and lower thermal breadths. This approximation has less than 10% relative error for TPCs that are not extremely skewed (*α* ∈ [0.06, 0.94]) and not too broad (*β* ≤ 0.5), which encompass the majority of TPC shapes that are likely to be encountered in practice (Figure S4). For large *β*, the interpretation of *β* as the upper thermal breadth at 88% of the peak height, as approximated in Equation 5, will no longer be accurate, but larger *β* always corresponds to broader TPCs, with the limit *β* → ∞ corresponding to a constant model where *r*(*T*) = *r_max_* in the [*T_min_*, *T_max_*] temperature range. Varying *α* and *β* makes it possible for flexTPC to describe unimodal curves with many different shapes (Figure 1, left panel).

An alternate parametrization of the flexTPC model that replaces *α* (the relative position of the thermal optimum) with the absolute optimum temperature *T_opt_* and *β* (the relative approximate upper thermal breadth) with the absolute approximate upper thermal breadth *B* = *β*(*T_max_* − *T_min_*) can also be constructed:

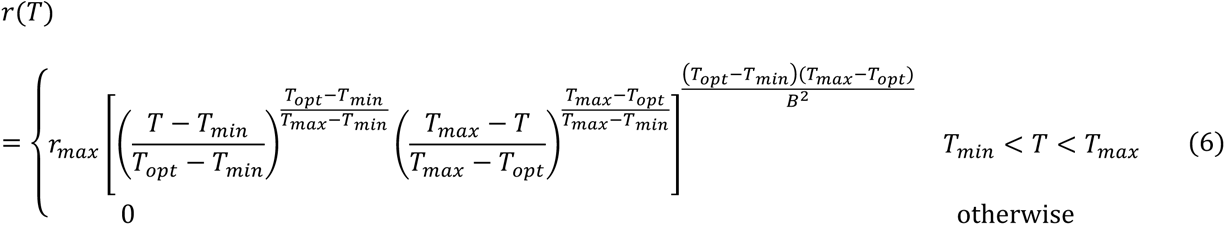

where *T_opt_* ∈ [*T_min_*, *T_max_*] and *B* > 0. In general, we expect Equation 6 to be useful for applied scientists who wish to automatically calculate confidence intervals on parameters of interest (absolute *T_opt_* and thermal breadth) using standard statistical software that performs nonlinear least squares or maximum likelihood estimation. Using this parametrization will lead to a confidence interval for *T_opt_* with no additional effort from the user of the statistical software.

However, there can be numerical issues with estimation for highly skewed curves where *T_opt_* is close to either *T_min_* or *T_max_*. When numerical issues arise, Equation 3 can be used instead.

Equation 3 is likely to be more useful when fitting TPCs through Bayesian methods, as it is more straightforward to provide a reasonable prior distribution for *α* (which lies in the interval from 0 to 1) than for *T_opt_* (which lies in-between two unknown model parameters: *T_min_* and *T_max_*). It is also simple to obtain posterior samples and credible intervals for *T_opt_* from MCMC output through Equation 4.

Equation 3 can also be used for maximum likelihood estimation: it is straightforward to obtain confidence intervals for *T_opt_*through bootstrap methods and any numerical issues regarding the optimal temperature “crossing-over” past the maximum or minimum temperatures can be avoided by constraining *α* to be in the unit interval. This parametrization also has the advantage of clearly separating the parameters that determine the shape (*α*, *β*) and location/scaling (*T_min_*, *T_max_*, *r_max_*) of the TPC.

### Datasets

To illustrate the predictive performance and applications of flexTPC, we compared it with the Briere model in various real-world datasets.

The `botrana` dataset consists of the developmental time of various life stages of the grapevine moth *Lobesia botrana* (eggs, instars 1-5, and pupae) measured at 14 temperatures, ranging from 8 to 34°C. This dataset, which was used to motivate development of the Briere models (Briere *et al*. 1999), was taken from Table 1 in (Briere & Pracros 1998). This dataset is expected to be one in which the Briere models perform well.

**Table 1.**
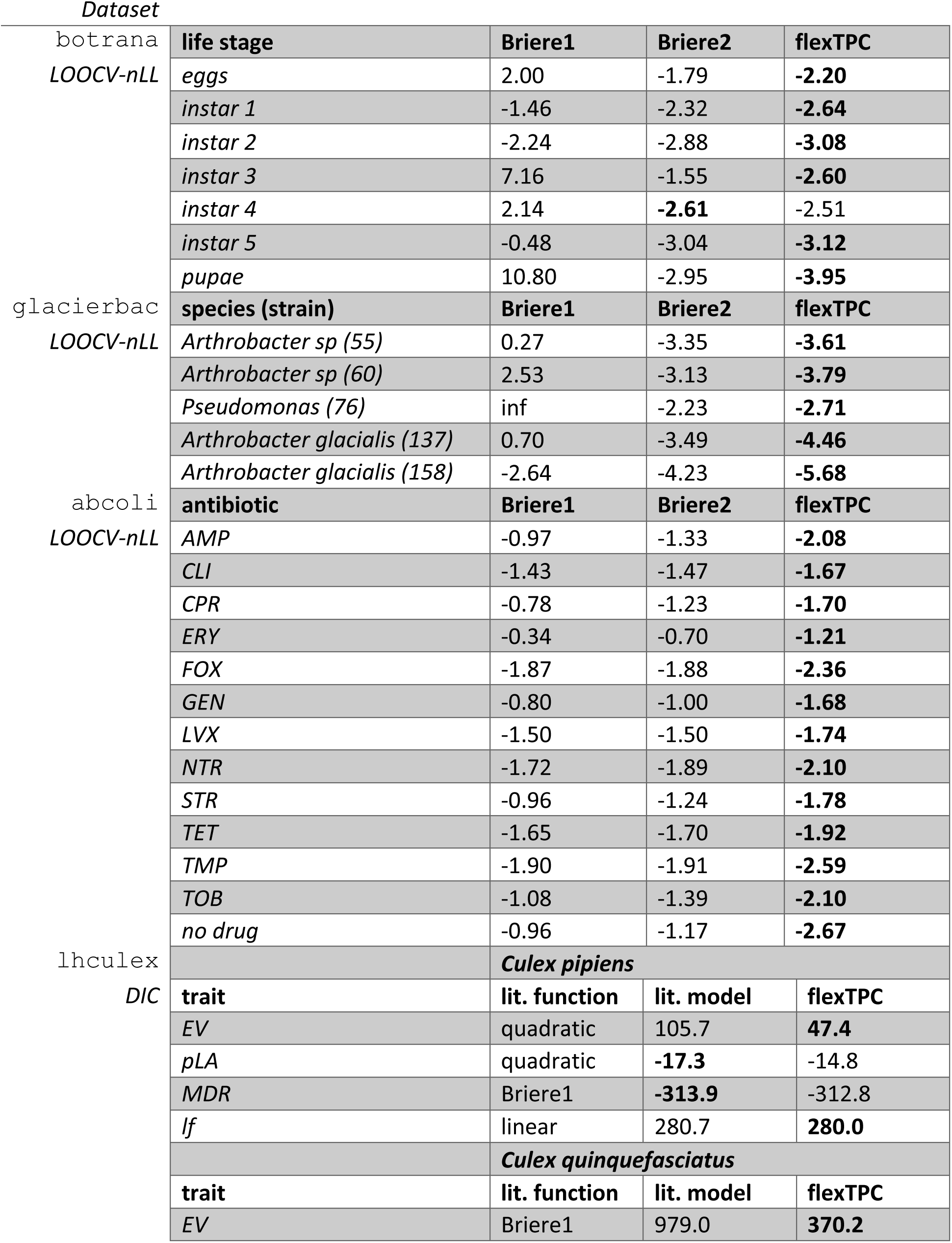
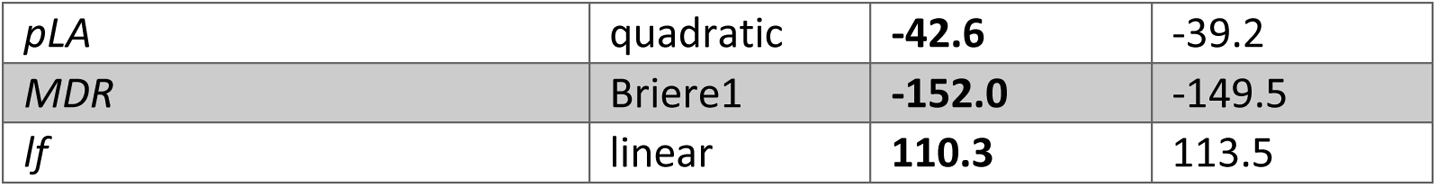
Model comparison in real-world datasets. We compare the predictive performance of flexTPC and Briere models. The best performing model has its values highlighted in bold. The model comparison criteria are indicated below the corresponding dataset. For datasets that were fit with a maximum likelihood approach (`botrana`, `glacierbac`, `lhculex`), we use mean leave one out cross-validated negative log-likelihood (LOOCV-nLL, lower is better) as the model comparison criterion to compare between the Briere1, Briere2, and flexTPC models. For the `lhculex` dataset, which was fit with a Bayesian approach, we use the Deviance Information Criterion (DIC, lower is better) as a model comparison criterion between a TPC functional form that was previously used in the literature to describe that trait (lit.function) and flexTPC.

The `glacierbac` dataset consists of the temperature dependence of the growth rate of bacterial *Arthrobacter* and *Pseudomonas* strains isolated from glacial deposits (Gounot 1976). This dataset was chosen to highlight the advantage of flexTPC over Briere in describing TPCs from organisms from cold environments.

The `abcoli` dataset (Cruz-Loya *et al*. 2021) consists of measurements of total growth after 24 hours of laboratory cultures of the bacterium *Escherichia coli* in the presence of various antibiotic backgrounds at seven temperatures. These antibiotics either kill or slow down the growth of *E. coli* in a temperature-dependent manner, modifying the shape of the TPC. This dataset was chosen to highlight the ability of flexTPC to describe curves of different shapes.

The `lhculex` dataset (Shocket *et al*. 2020) corresponds to various mosquito temperature-dependent life history traits (egg viability, probability of larval survival to adulthood, development rate, and female adult lifespan) from *Culex pipiens* and *Culex quinquefasciatus*. These traits have been previously modeled with different functional forms (linear, quadratic, and Briere1). This dataset was chosen to highlight the ability of flexTPC to fit curves of various shapes for which different functional forms were previously needed.

### Parameter estimation

A nonlinear regression approach was used to fit the Briere1, Briere2, and flexTPC models to the `botrana, glacierbac,` and `abcoli` datasets through maximum likelihood estimation. The following model was used for the `botrana` and `glacierbac` datasets:

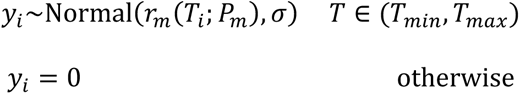

where *y_i_* is the observed response at temperature *T_i_*, *σ* the standard deviation of the data, *r_m_* the temperature response curve model (either Briere1, Briere2, or flexTPC), and *P_i_* the set of all parameters from the corresponding TPC model being fit. For example, *P*_Briere1_ = {*T_min_*, *T_max_*, *c*}.

In the `abcoli` dataset, the response variable is optical density, which does not have zero values. For this dataset the model used was:

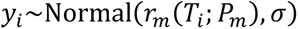

As a criterion for model selection, we compared the negative log-likelihood obtained under leave one out cross-validation (LOOCV-nLL) for all models in the datasets described above. This is a measure of the predictive out-of-sample model performance that is asymptotically equivalent to AIC (Stone 1977) but makes fewer assumptions, and has been recommended as the approach of choice for model selection when computationally feasible (Yates *et al*. 2023). It consists of removing each data point in turn, fitting the model with maximum likelihood on the remaining data points, and evaluating the negative log-likelihood (nLL) in the removed data point (which is a measure of the quality of the model prediction for a data point that was not used in fitting). We report the mean nLL when each data point is removed in turn. Alternate model comparison criteria (AIC and BIC) are reported in the Supplemental Information.

### Bayesian parameter estimation for mosquito trait data

For the `lhculex` dataset, we followed a Bayesian approach for parameter estimation. This makes it possible to fit curves with reasonable thermal limits for traits that lack data at low temperatures using weakly informative prior distributions and illustrates the benefits of fitting flexTPC in a Bayesian context. For each mosquito life history trait, flexTPC was compared to a TPC functional form that was used previously to describe the data being modeled, which varied by trait (Shocket *et al*. 2020). Deviance Information Criterion (DIC) (Spiegelhalter *et al*. 2002) was used as a model selection criterion. For more details, see Table 1 and the Supplemental Information.

Models were fit using Markov Chain Monte Carlo (MCMC) with the r2jags R package, an interface for JAGS (Just Another Gibbs Sampler) (Plummer 2003). Four independent MCMC chains were run for 300,000 iterations, discarding the first 50,000 iterations as burn-in. The resulting MCMC chains were thinned, saving every eight iterations. Chain convergence was monitored both by visual inspection of trace plots and density plots of the individual chains and by ensuring the potential scale reduction factor *R̂* < 1.01 for all parameters.

## Results

In this work, we present flexTPC—a flexible model for unimodal thermal performance curves (TPCs) in which the optimum temperature can lie at any point in between the minimum and maximum temperatures. This model is parameterized in terms of biologically meaningful quantities and can describe TPCs of a wide variety of shapes (Figure 1). We compare the performance of flexTPC to that of the Briere1 and Briere2 models (Equation 1), which are phenomenological models for TPCs that are popular in applications in various real-world datasets.

### Insect developmental rates

The Briere models were initially developed to describe the thermal dependence of insect developmental rates. We compared the flexTPC and Briere models for describing Briere and Pacros’s data on the rates of development of the life stages of the grapevine moth *Lobesia botrana* (Figures 2 (left panel), S1) to evaluate the relative performance of these models in a real dataset for which the Briere models would be typically used.

**Figure 2.**
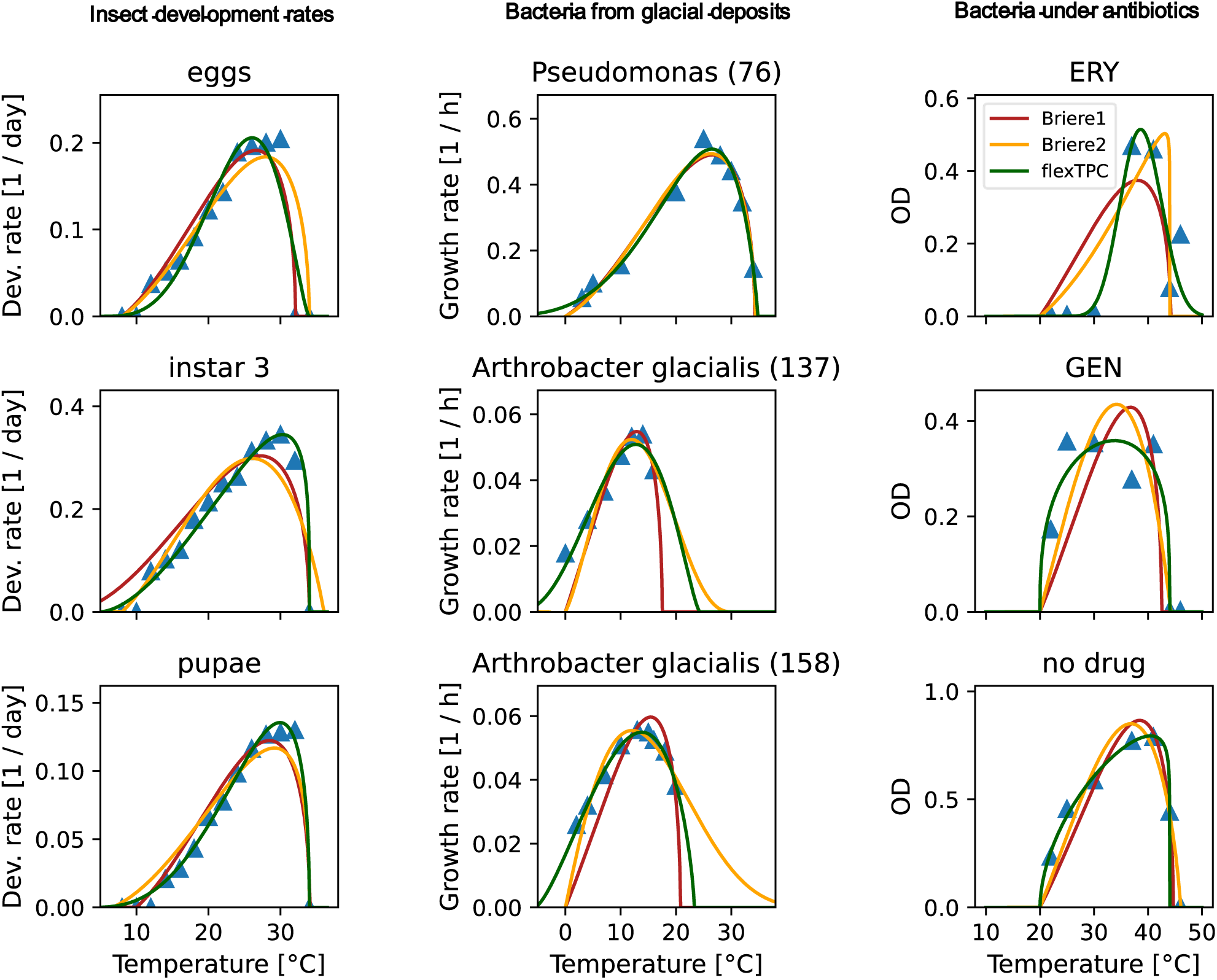
FlexTPC outperforms the Briere1 and Briere2 models in various real-world datasets. Data (shown as blue triangles) and fitted TPC models (Briere1: red lines, Briere2: yellow lines, flexTPC: green lines) for selected examples from various real-world datasets (`botrana`, `glacierbac` and `abcoli`, see Methods). *Left column.* Rate of development of various life stages of the grapevine moth *Lobesia botrana*. A subset of the life stages (eggs, instar 3 and pupae) is shown. *Middle column.* Growth rate of psychrophile bacterial species (*Pseudomonas* and *Arthrobacter glacialis*) isolated from glacial deposits. *Right column.* Optical density (OD, a proxy for the number of bacteria) of *Escherichia coli* cultures after 24-hour growth under various antibiotic backgrounds (ERY: erythromycin, GEN: gentamycin, no drug: growth media without antibiotics). The fitted TPC models for all traits in each dataset are shown in Figures S1-S3 in the Supplemental Information.

Based on leave-one-out cross validation (LOOCV), we found that flexTPC was the best performing model for six life stages (eggs, instars 1, 2, 3, 5 and pupae) while the Briere2 model was the best performing model for one life stage (instar 4; Table 1). The Briere1 model was the worst performing model for all life stages in this dataset.

Organisms that live below freezing temperatures

The Briere models force trait performance to be zero at *T* = 0°C and are thus unable to describe thermal performance curves for traits of living organisms that function below freezing temperatures. In order to provide a real-world example, we next compared the Briere and flexTPC models for describing the growth rate of three facultative psychrophile bacterial strains (*Arthrobacter sp* strain SI 55, *Arthrobacter sp* strain SI 60, and *Pseudomonas* strain SII 76) and two obligate psychrophile strains (*Arthrobacter glacialis* strains SI 137 and SI 158) isolated from glaciers (Gounot 1976) (Figures 2 (middle column), S2). We found that flexTPC provides better fits than both Briere models for all bacterial species in the dataset (Table 1). This was especially so for both *Arthrobacter glacialis* strains since they exhibit substantial growth at and below 0°C, which is impossible to capture with the Briere models.

Thermal performance curves (TPCs) of varying shapes

Thermal performance curves (especially those for growth and developmental rates) are often left-skewed, with the temperature optimum closer to the maximum than the minimum temperature for the trait. However, some traits have symmetric or right-skewed TPCs, and environmental stressors can change the shape of TPCs (Bestion *et al*. 2018; Brett *et al*. 1969; Cruz-Loya *et al*. 2021; Cuppers *et al*. 1997). As a real-world example, we next considered a dataset consisting of the temperature-dependent growth of *Escherichia coli* under 12 different antibiotics, and a control condition in the absence of antibiotics (Cruz-Loya *et al*. 2021). We again compared the fit of the Briere1, Briere2, and flexTPC models (Figures 2 (right column), S3, Table 1).

While the TPC of *E. coli* growth is left-skewed in the absence of antibiotics, its shape can be modified in their presence because antibiotic effectiveness can vary at different temperatures. Some antibiotics give rise to left-skewed curves (e.g., TET, TMP, FOX), while others result in curves that are closer to symmetric and can be either nearly flat (GEN, TOB, STR) or narrow (ERY). FlexTPC was the best performing model for all 13 antibiotic backgrounds in this dataset (Table 1) and is the only model out of the three that can describe TPCs that are symmetric or right-skewed.

Fitting thermal performance curves that vary in shape across multiple traits and species Organisms have multiple temperature dependent traits, giving rise to TPCs that can have different shapes. In practice, this has often meant that a different TPC functional form (such as Briere or quadratic) must be chosen for each trait, and sometimes even for the same trait in different species. This raises the issue that the inferred parameters (like minimum, optimal, and maximum temperatures) may differ across traits or species partially because of using different functional forms rather than only because of the data. A flexible model such as flexTPC makes it possible to compare TPCs of different shapes with the same model, allowing the direct comparison of inferred parameters. In addition, having interpretable model parameters allows the use of informative Bayesian priors based on curves fit to related species or knowledge of the temperature range in the habitat of the species of interest.

As an example, we fit TPC models to a dataset with four life history traits (lifespan, egg viability, larval survival to adulthood, and mosquito development rate) of the mosquitoes *Culex pipiens* and *Culex quinquefasciatus* using a Bayesian approach (Figure 3). In a previous study (Shocket *et al*. 2020), these data were analyzed using various different functional forms (linear, quadratic, and Briere), depending on the trait and species (Table 1). We find that flexTPC gives very similar fits to using these different models for lifespan, larval survival, and development rate. Moreover, it provides substantially better fits for egg viability compared to the previous models chosen in the literature (quadratic for *Cx. pipiens* and Briere1 for *Cx. quinquefasciatus*).

**Figure 3.**
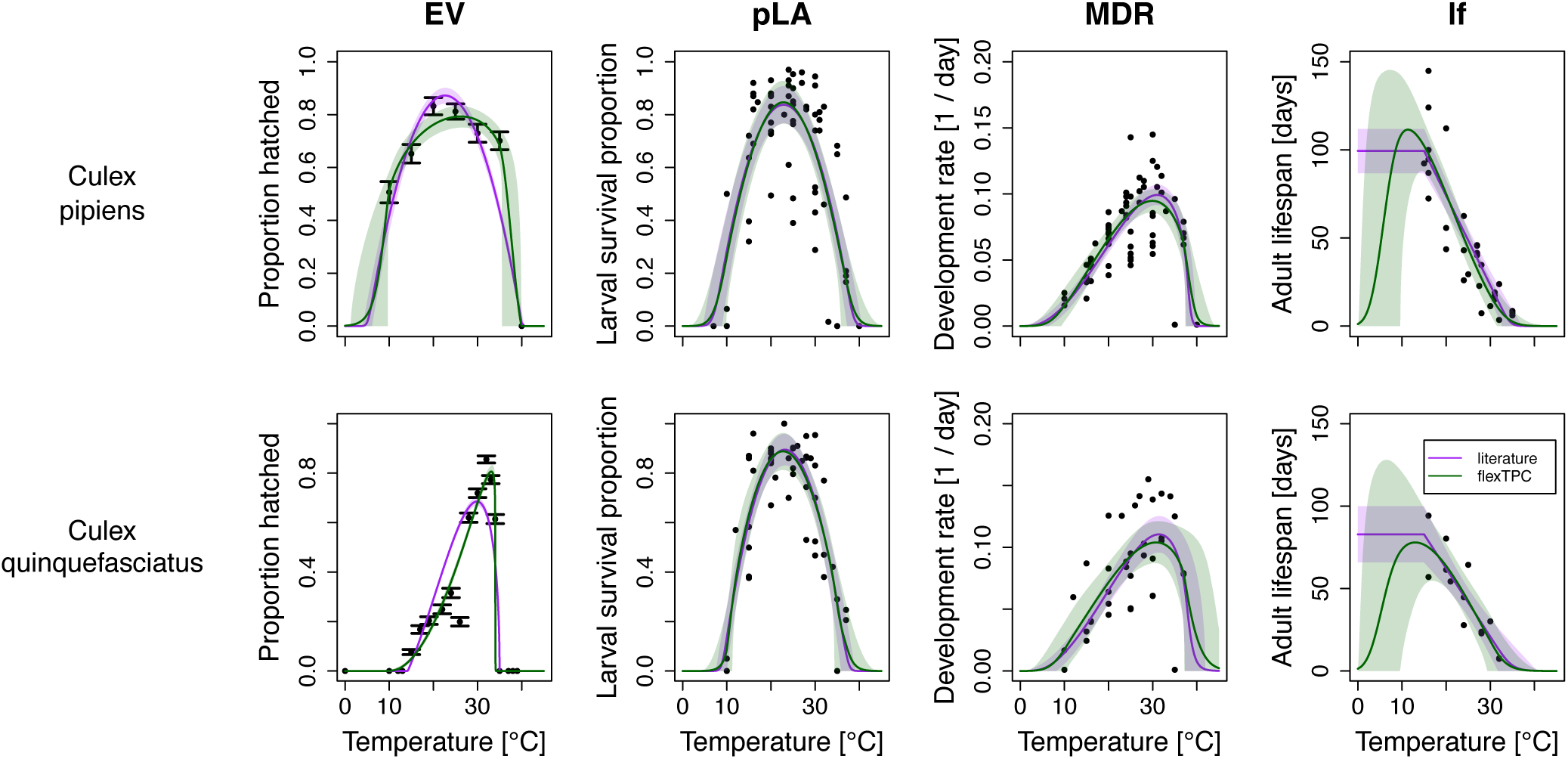
FlexTPC can be used to fit the thermal performance of multiple traits with different shapes that typically require the choice of different TPC models. We show Bayesian fits to egg viability (EV), probability of larval survival to adulthood (pLA), mosquito development rate (MDR), and female adult lifespan (lf) for *Culex pipiens* (top row) and *Culex quinquefasciatus* mosquitoes (bottom row). These traits have very different shapes and different TPC models have been used in the past to fit data from these traits. We compare the flexTPC model (green) with a previously used TPC functional form that varies by trait and species (purple, see Table 4 for the specific model for each trait). Lines correspond to posterior means and shaded regions to 95% credible intervals, which represent the uncertainty of the true value of the TPC at each temperature.

For adult lifespan, flexTPC results in a near-identical fit to that of a piecewise linear model (which was previously used to describe this trait) within the range of the data. Although this dataset does not contain temperatures low enough to observe a reduction in lifespan, it must necessarily decrease at lower temperatures, so it is likely more realistic to model this trait as a right-skewed unimodal TPC (as can be done with flexTPC) rather than a linear model. If Bayesian methods are used, this can be done even in cases where there is a lack of data near temperature extremes.

In Bayesian approaches, uncertainty in model parameters is described by probability distributions. Before the analysis, a prior distribution for each parameter is chosen that represents how likely each parameter value is assumed to be *a priori* (before observing the data). Prior distributions can be based on biological knowledge from previous experiments in related species or known characteristics of the habitat of the population being studied. For example, as the mosquito species of interest are ectotherms that live in temperate (*Cx. pipiens*) or tropical/subtropical (*Cx. quinquefasciatus)* climates, we assume that *T_min_* and *T_max_* for adult lifespan are *a priori* 95% likely to be in the interval (0°C, 10°C) and (25°C, 45°C), respectively. Choosing reasonable prior distributions based on biological knowledge is much easier when the model parameters are interpretable (e.g., for minimum and maximum temperatures and the maximum trait value) rather than mathematical constants with no direct biological meaning.

Because of its interpretable parameters (Figure 4 and Box 1), flexTPC is well-suited for Bayesian parameter estimation.

**Figure 4.**
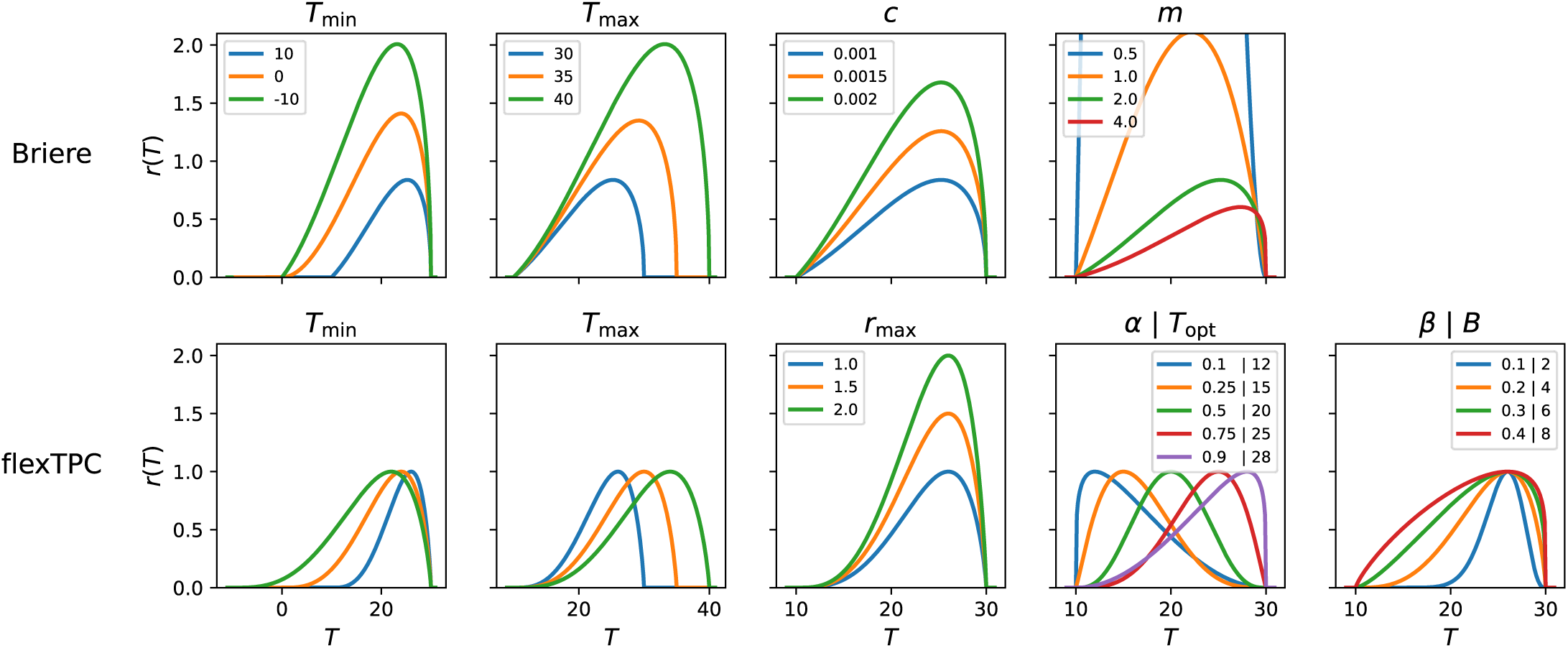
Single parameter changes in the Briere and flexTPC models. In each panel, we show the effects on the thermal performance curve when a single parameter of the corresponding TPC model is changed while keeping all other parameters constant. For parameters other than *m* in the Briere model, a fixed value of *m* = 2 is used (corresponding to the Briere1 model). We show the parameter values for both parametrizations of flexTPC (Equations 3 and 6), which differ on whether the optimal temperature and approximate upper thermal breadth are in unitless (*α*, *β*) or dimensional (*T_opt_*, *B*) form, but are otherwise identical and describe the same set of curves. Since flexTPC has biologically interpretable parameters, changing a single parameter (e.g., *T_min_*) will change the thermal performance curve in a predictable way (as the rest of the parameters that are kept constant correspond to known curve properties). In contrast, in a model where some parameters are mathematical constants without a direct biological interpretation, changing a parameter can lead to unintuitive and possibly unintended changes in the thermal performance curve (e.g., changing *T_min_* also leads to changes on the height of the curve for the Briere model). This has important consequences when modeling changes in TPCs due to evolutionary or environmental factors, and when interpreting sensitivity analyses of derived quantities from TPC models (see Box 1). Note that decreasing parameter *T_min_* to negative values in the Briere model does not lead to models with positive performance below 0°C (see Methods).

#### Box 1

**Advantages of thermal performance curve models with biologically interpretable parameters**

For many applications (for example, studying the evolution of TPCs or predicting the effect of thermal adaptation on infectious disease spread), it is of interest to model how thermal performance curves change across time, across space, in the presence of a stressor other than temperature, and/or when exposed to other factors that vary across populations. It is natural to do this by making assumptions about how parameters of interest (e.g., minimum, optimum, or maximum temperatures) change as a function of the variable of interest. However, when some parameters in the chosen TPC functional form are mathematical constants without a clear biological interpretation, this can lead to unintuitive changes in the predicted values for the TPC, even when the parameter being modified is interpretable.

To illustrate this, we show the effects of changing a single parameter while keeping all other parameters constant for the Briere and flexTPC models (Figure 4). In the Briere model there is a multiplicative constant *c* that is proportional to the height of the curve when all other model parameters are fixed. Changing the value of *c* while keeping the other model parameters constant will change the TPC in a predictable way by modifying its height while keeping the same minimum and maximum temperatures. However, changing the value of a different model parameter in the Briere model (e.g., *T_min_* or *T_max_*, which are interpretable parameters) while keeping all other parameters constant will not keep the height of the curve constant, as the value of *c* that is needed to keep the same height changes when the other model parameters change. In contrast, in the flexTPC model the maximum trait value *r_max_* (i.e., the curve height) is explicitly a model parameter. Thus, keeping *r_max_* constant will keep the same TPC height regardless of the values of the other parameters. When modeling changes in TPCs, it is advantageous to choose a functional form where parameters are biologically interpretable, especially if it is of interest to assume certain aspects of the TPC remain constant or change in a predictable way. This will lead to a clearer interpretation of changes in model parameters which is not confounded by changes in other aspects of the TPC that are not of interest.

Using TPC models where some of the parameters are mathematical constants without a biological interpretation can lead to potentially misleading conclusions in applications that require the interpretation of partial derivatives of the model or quantities derived from them. Importantly, this includes sensitivity analyses of mathematical models that include TPCs as a submodel (such as infectious disease or predator-prey models) with respect to the underlying parameters of the TPC functional form. For example, sensitivity analysis based on partial derivatives might indicate that the transmission of a disease is very sensitive to the parameter *T_max_* of a TPC modeled with the Briere1 function. However, as increasing *T_max_* (while keeping all other parameters constant) also increases the height of the TPC, this could be either due to the increased maximum temperature or the increased curve height. In contrast, using a model where all parameters have a clear biological interpretation (and where the maximum value of the TPC is an explicit parameter) enables separating the effect of increasing the maximum temperature and increasing the curve height.

In general, parametrizing models in terms of biologically interpretable quantities is useful as it makes it possible to keep them constant or to change them in specified ways when varying other parameters (as needed for modeling change in TPCs). It is also advisable to explore the effects of changing individual parameters in the TPCs to be aware of what aspects of the curve are being modified by the parameter in question when interpreting sensitivity analyses.

## Discussion

In this work we introduce flexTPC, a flexible mathematical model for thermal performance curves (TPCs) that can describe unimodal TPCs of various shapes (including left-skewed, symmetric, and right-skewed curves). FlexTPC is mathematically equivalent to the Beta model (Yin *et al*. 1995), but is reparametrized to be biologically interpretable and better suited for applied ecology and infectious disease applications. We show that this model addresses various limitations of the Briere models, such as not being able to describe TPCs from species that can survive below freezing temperatures, or TPCs that vary in skewness/thermal breadth. This leads to better predictive performance in various real-world datasets. Based on these results, we propose flexTPC as a general-purpose descriptive model to describe unimodal TPCs.

FlexTPC is parametrized in terms of biologically meaningful quantities that are of interest to ecologists: the minimum and maximum temperatures, the maximum value of the trait, a choice of either the relative or absolute position of the optimum temperature, and a choice of the approximate relative or absolute upper thermal breadth. This has several advantages when compared to models in which some parameters are mathematical constants without a clear interpretation. First, the model behaves more predictably when changing its parameter values, since these quantities can be kept constant or modified intentionally as opposed to changing in possibly unintuitive ways as other parameters vary (Figure 4). This aids in the clear interpretation of parameter sensitivity analysis and facilitates modeling how TPCs change over time and/or space (Box 1). Second, it simplifies finding reasonable initial values for the parameters when fitting the model with optimization-based methods (e.g., least squares or maximum likelihood estimation). Third, statistics such as confidence intervals can often be obtained automatically with standard software when a quantity of interest is an explicit parameter of the model. Lastly, an interpretable parametrization makes it easier to incorporate information from previous experiments in similar species or other sources (e.g., the environmental temperature range from the habitat of the organism) when using informative priors in Bayesian approaches to parameter inference.

FlexTPC has several important advantages over other popular models like the Briere models. First, in any model describing TPCs, at least three parameters are necessary in order to set the curve height and the minimum and maximum temperatures independently. Because of this, the optimal temperature in any TPC model that has three parameters or fewer (like the Briere1 model or the quadratic model) will necessarily be a deterministic function of some subset of these parameters. This may lead to biased estimates for the optimum temperature (and the other parameters involved in the deterministic relationship) whenever the true relationship between these parameters deviates from the implicit assumption made by the TPC functional form, which often lacks a biological justification in phenomenological models. FlexTPC (and Briere2) can vary the optimum temperature for fixed values of the minimum and maximum temperature and are thus likely better suited for estimating optimal temperatures, especially when thermal limits are tightly constrained by the data. Conversely, when using the Briere1 function to describe a TPC where most data are near the optimum, the estimated thermal minimum and maximum might be inaccurate due to the constraints imposed by the functional form.

Second, organisms may function below freezing temperatures, and while the Briere1 and the Briere2 models cannot describe positive performance below freezing, flexTPC can describe TPCs at any temperature range (Figures 2, S2). Although it is possible to use the Briere models in these cases by shifting the model in the temperature axis, this requires choosing an arbitrary temperature shift, and the shape of the resulting TPC depends on the chosen shift (Figure S5).

Another advantage of flexTPC over the Briere models is its ability to describe TPCs of many different shapes. This will be especially useful in studies comparing multiple TPCs from different traits and/or from different organisms. Currently, different functional forms are commonly used in these studies when the TPC shape changes across species or traits. This can potentially introduce issues when comparing inferred parameters, as parameters might vary between conditions partially due to the use of a different model rather than because of meaningful differences in the data. This issue can be avoided by using a flexible model that allows fitting all conditions with the same functional form.

As flexTPC is a more complex model that the Briere models, with five free parameters, it is natural to consider whether it can be used in data-limited situations where measurements are only available at a few temperatures, as frequently occurs in lab and field data. In this work we show that, despite this additional complexity, flexTPC has better predictive performance than the Briere1 and Briere2 models in many real-world scenarios. Moreover, as illustrated in the data for mosquito lifespan (Figure 3), flexTPC can be used in situations with limited data at some temperature ranges when using Bayesian methods. Even in cases with severe data limitations, the use of a flexible model with Bayesian methods with strongly informative priors based on biological knowledge of the species being modeled and its habitat may be preferable to the use of a more parsimonious model that assumes a strong relationship between the optimal, minimum, and maximum temperatures without biological justification, especially when the main purpose of the analysis is to estimate an optimal temperature. However, more parsimonious models can be obtained from the flexTPC equation for researchers under severe data constraints that do not wish to take a Bayesian approach to parameter inference (see Supplemental Information).

Our work shows that flexTPC is a general-purpose model for unimodal TPCs that is well-suited for comparing populations or experimental conditions where the curves may vary in thermal breadth and skewness. To our knowledge, flexTPC is the first descriptive TPC model to simultaneously have an explicit parameter corresponding to all of the main TPC features of interest for ecologists—the temperature minimum, maximum, and optimum, along with the maximum trait performance value and thermal breadth. This inclusion of parameters of interest results in a model that is both flexible and interpretable, which we believe will be useful for both fitting empirical data and for theoretical work that models how TPCs change under evolution or in the presence of external factors like other stressors. FlexTPC can also be used as a flexible functional form to describe the response of biological traits to other environmental factors (e.g., precipitation or humidity) when these responses are unimodal.

## Acknowledgements

MCL and EAM were supported by NIH grants R35GM133439 and R01AI168097. MCL was also supported by a Stanford PRISM Baker Fellowship. EAM was also supported by NSF grant DEB-2011147 with Fogarty International Center, NIH grant R01AI102918, and seed grants from the Stanford Woods Institute for the Environment, King Center on Global Development, and Center for Innovation in Global Health. VMS was supported by a James F. McDonnell Foundation Complex Systems Scholar Award.

## Data and code availability

All data and code are provided in a GitHub repository.

