## Supplemental Information for "A flexible model for thermal performance curves"

#### Supplemental Figures

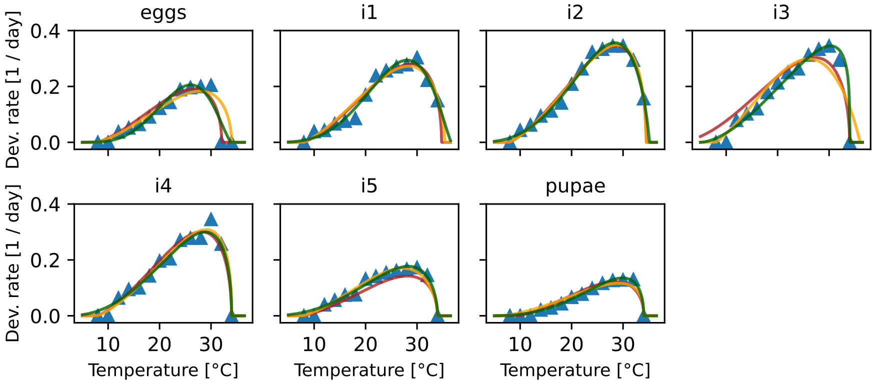


Figure S1: *FlexTPC has similar or better performance than the Briere models when describing insect developmental rates.* The Briere1, Briere2 and flexTPC models are compared in the `botrana` dataset. Experimental measurements of the developmental rate (defined as the inverse of mean time to progression) of various life stages (eggs, instars 1 through 5 and pupae) of the grapevine moth *Lobesia botrana* at various temperatures are shown as blue triangles. The fitted models are shown as lines (Briere1: red, Briere2: yellow, flexTPC: green).

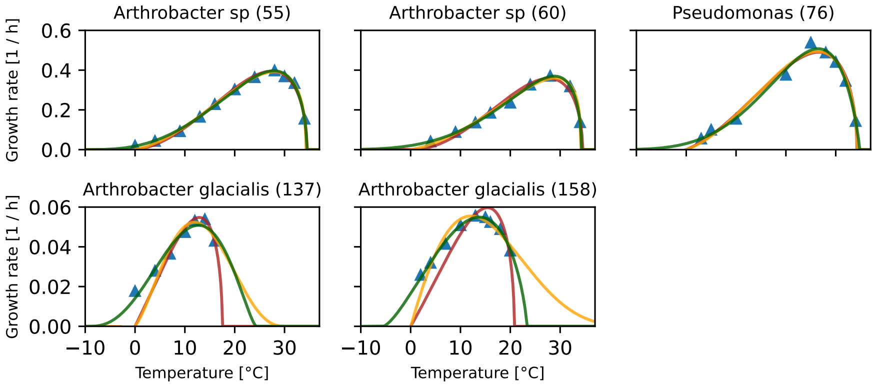


Figure S2: *FlexTPC outperforms the Briere models when describing TPCs of organisms from cold environments.* The Briere1, Briere2 and flexTPC models are compared in the `glacierbac` dataset. Experimental measurements of the growth rate of five different bacterial strains are shown as blue triangles. The fitted models are shown as lines (Briere1: red, Briere2: yellow, flexTPC: green). The strains shown in the top row are facultative psychrophiles while the strains shown in the bottom are obligate psychrophiles. Due to their mathematical structure, the Briere models cannot describe TPCs for which the minimum temperature is less than 0°C.

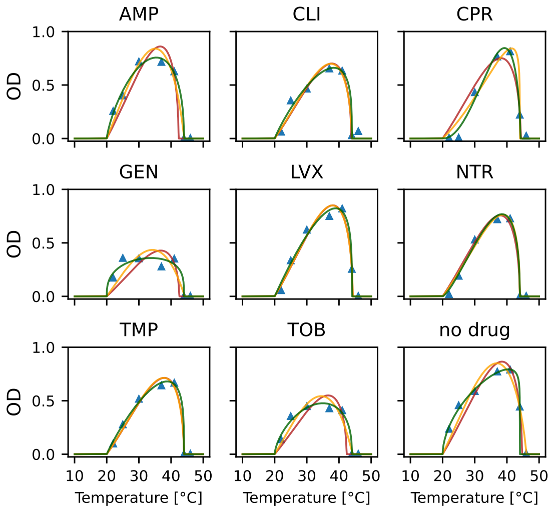


Figure S3: *FlexTPC can describe thermal performance curves of various shapes that arise under stressors.* We show the model fits of flexTPC and the Briere models in the `abcoli` dataset. The optical density (a proxy for the total amount of bacteria) of an *E. coli* culture after 24 hours of growth under various antibiotic backgrounds (AMP: ampicillin, CLI: clindamycin, CPR: ciprofloxacin, ERY: erythromycin, FOX: cefoxitin, GEN: gentamycin, LVX: levofloxacin, NTR: nitrofurantoin, STR: streptomycin, TET: tetracyclin, TOB: tobramycin, no drug: no antibiotic) is shown as blue triangles. The fitted models are shown as lines (Briere1: red, Briere2: yellow, flexTPC: green). Note that the presence of antibiotics changes the shape of the thermal performance curve.

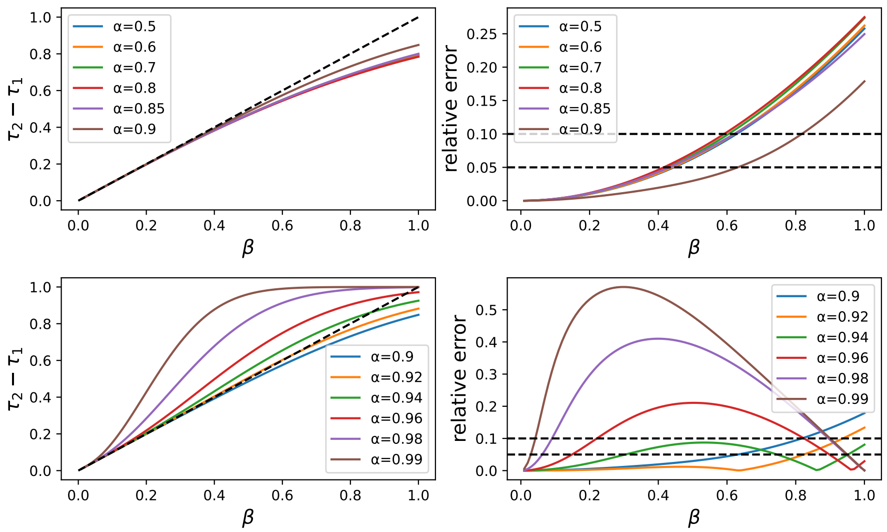


Figure S4: *Accuracy of upper thermal breadth approximation.* Parameter $\beta$ of the flexTPC model approximates the ratio between the upper thermal breadth $T_2 - T_1$ (defined as the width of the temperature interval for which $\frac{r(T)}{r_{\max}} > e^{-\frac{1}{8}} \approx 0.88$) and the lower thermal breadth $T_{\max} - T_{\min}$ (defined as the width of the temperature interval for which $\frac{r(T)}{r_{\max}} > 0$). *Left.* The model parameter $\beta$ is compared to the true thermal breadth ratio $\tau_2 - \tau_1 = \frac{T_2 - T_1}{T_{\max} - T_{\min}}$ (where $\tau_2 - \tau_1$ was obtained numerically) for various values of parameter $\alpha$, which determines the location of the thermal optimum relative to $T_{\min}$ and $T_{\max}$. The identity line is shown as a dotted line for comparison. We only show symmetric and left-skewed curves (i.e. when $\alpha \geq 0.5$) since, by symmetry, the error behavior will be the same for the corresponding right-skewed curve obtained by replacing $\alpha$with $1 - \alpha$. *Right.* The relative error $\left|\frac{\beta - (\tau_2 - \tau_1)}{\tau_2 - \tau_1}\right|$ of $\beta$ when approximating $\tau_2 - \tau_1$ is shown for various values of $\alpha$. Dotted lines show relative errors of 5% and 10%.

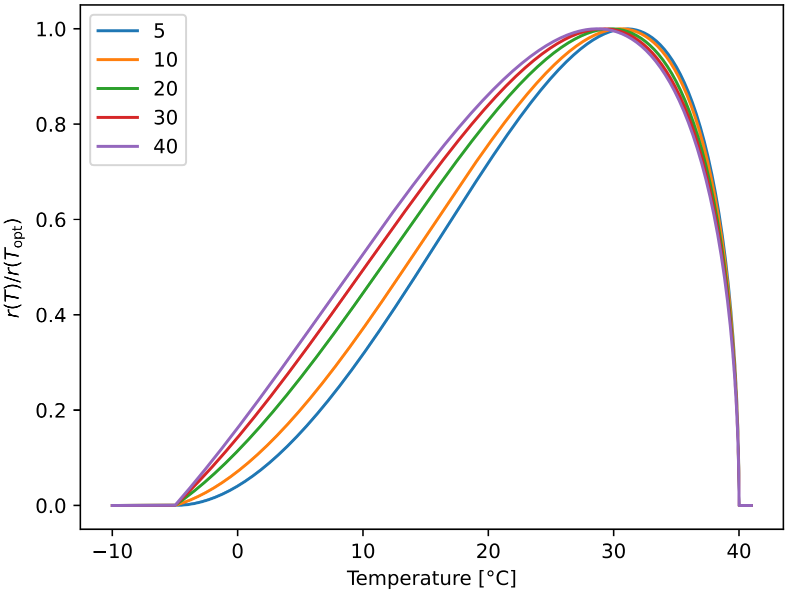


Figure S5: *Shifing the Briere model to accomodate traits with nonzero peformance at lower temperatures changes the shape of the curve.* An attempt to use the Briere model below freezing temperatures could be to shift it left by an offset $T_{\mathrm{offset}}$ (see shifted Briere model section in the Supplemental Information). We show the curve shapes (normalized by the maximum value to more easily compare the curve shapes) of different choices of $T_{\mathrm{offset}}$ while keeping all other model parameters constant. The resulting TPCs can describe TPCs below freezing temperatures, but the curve shape depends on the choice of offset temperature, which is effectively an additional model parameter.

### Mathematical derivation of the flexTPC model

We start with the equation for the four-parameter Briere2 model for thermal performance curves

$$r(T) = \begin{cases} cT(T - T_{\min})(T_{\max} - T)^{\frac{1}{m}} & T_{\min} < T < T_{\max} \\ 0 & \text{otherwise} \end{cases}$$

and modify it as follows:

$$r(T) = \begin{cases} c(T - T_{\min})^{a}(T_{\max} - T)^{b} & T_{\min} < T < T_{\max} \\ 0 & \text{otherwise} \end{cases} \tag{S1}$$

In doing this we change the model in two important ways. First, we remove the root at zero: this will make the model suitable for describing TPCs of organisms that function below freezing temperatures. Second, we introduce an exponent $a$ for the first factor to make the model more flexible and replace the power $1/m$ by a constant $b$. This leads to a flexible model that can describe temperature performance curves of many different shapes that is equivalent to the Beta model [2] if we replace the constant $c$ with a different constant $k$ by taking $c = e^k$.

However, this form of the model contains three arbitrary mathematical constants $a, b, c$ without a clear biological meaning, making it difficult to interpret the model parameters, determine initial values for curve fitting and set reasonable prior distributions with Bayesian approaches. To make this model more useful for ecological applications, we reparametrize the model by replacing these arbitrary constants with biologically meaningful quantities. We start by finding the optimal temperature. If we take natural logarithms and differentiate we have

$$\begin{aligned} \frac{d}{dT}\ln r &= \frac{d}{dT}\left[\ln c + a\ln(T - T_{\min}) + b\ln(T_{\max} - T)\right] \\ &= \frac{a}{T - T_{\min}} - \frac{b}{T_{\max} - T} \end{aligned}$$

By Fermat's theorem on extrema, $\ln r(T)$ is maximized when $\frac{d}{dT}\ln r = 0$. Since the natural logarithm is a monotonically increasing function, this condition also maximizes $r(T)$. Thus, we can find the optimum temperature $T_{\text{opt}}$ by solving the equation

$$0 = \frac{a}{T_{\text{opt}} - T_{\min}} - \frac{b}{T_{\max} - T_{\text{opt}}}$$

which yields

$$T_{\text{opt}} = \alpha T_{\max} + (1 - \alpha)T_{\min} \tag{S2}$$

where $\alpha = \frac{a}{a+b}$. Rearranging this equation yields $\alpha = \frac{T_{\text{opt}} - T_{\min}}{T_{\max} - T_{\min}}$, which shows that $\alpha$ can be interpreted as a rescaled (and dimensionless) optimal temperature. In other words, it represents the position of the optimal temperature relative to the minimum and maximum, with the extreme values $\alpha = 0$ corresponding to $T_{\mathrm{opt}} = T_{\mathrm{min}}$ and $\alpha = 1$ to $T_{\mathrm{opt}} = T_{\mathrm{max}}$.

If we take $\alpha = \frac{a}{a+b}$ and define $s = a+b$, we have that $a = \alpha s$ and $b = (1-\alpha)s$. Substituting these constants in Equation S1 yields

$$r(T) = \begin{cases} c(T - T_{\mathrm{min}})^{\alpha s}(T_{\mathrm{max}} - T)^{(1-\alpha)s} & T_{\mathrm{min}} < T < T_{\mathrm{max}} \\ 0 & \text{otherwise} \end{cases} \tag{S3}$$

Now, we replace the arbitrary constant $c$ with the maximum trait value $r_{\mathrm{max}}$ by noting that by definition $r_{\mathrm{max}} = r(T_{\mathrm{opt}})$, which gives the following equation

$$r_{\mathrm{max}} = c(T_{\mathrm{opt}} - T_{\mathrm{min}})^{\alpha s}(T_{\mathrm{max}} - T_{\mathrm{opt}})^{(1-\alpha)s}$$

Substituting Equation S2 for $T_{\mathrm{opt}}$ into this expression and solving for $c$ yields

$$c = \frac{r_{\mathrm{max}}}{\left[\alpha^{\alpha}(1-\alpha)^{1-\alpha}(T_{\mathrm{max}} - T_{\mathrm{min}})\right]^{s}}$$

Substituting this expression for $c$ into Equation S3 and rearranging the resulting expression yields the following:

$$r(T) = \begin{cases} r_{\mathrm{max}} \left[ \left(\frac{T - T_{\mathrm{min}}}{\alpha}\right)^{\alpha} \left(\frac{T_{\mathrm{max}} - T}{1-\alpha}\right)^{1-\alpha} \left(\frac{1}{T_{\mathrm{max}} - T_{\mathrm{min}}}\right) \right]^{s} & T_{\mathrm{min}} < T < T_{\mathrm{max}} \\ 0 & \text{otherwise} \end{cases} \tag{S4}$$

This version of the model was previously derived in [1], where it is referred to as the modified Briere model. An alternate parametrization that has the optimum temperature $T_{\mathrm{opt}} \in [T_{\mathrm{min}}, T_{\mathrm{max}}]$ explicitly can be found by substituting the expression $\alpha = \frac{T_{\mathrm{opt}} - T_{\mathrm{min}}}{T_{\mathrm{max}} - T_{\mathrm{min}}}$ and rearranging to get

$$r(T) = \begin{cases} r_{\mathrm{max}} \left[ \left(\frac{T - T_{\mathrm{min}}}{T_{\mathrm{opt}} - T_{\mathrm{min}}}\right)^{\frac{T_{\mathrm{opt}} - T_{\mathrm{min}}}{T_{\mathrm{max}} - T_{\mathrm{min}}}} \left(\frac{T_{\mathrm{max}} - T}{T_{\mathrm{max}} - T_{\mathrm{opt}}}\right)^{\frac{T_{\mathrm{max}} - T_{\mathrm{opt}}}{T_{\mathrm{max}} - T_{\mathrm{min}}}} \right]^{s} & T_{\mathrm{min}} < T < T_{\mathrm{max}} \\ 0 & \text{otherwise} \end{cases} \tag{S5}$$

We can find the parameters of the model that determine its shape by nondimensionalizing the model, which will remove any parameters that determine the location and scaling of the thermal performance curve without changing its shape. To do this, we can define a nondimensional temperature $\tau = \frac{T - T_{\mathrm{min}}}{T_{\mathrm{max}} - T_{\mathrm{min}}}$ (where $\tau$ and $T$ have a relationship that is analogous to that between $\alpha$ and $T_{\mathrm{opt}}$) and a nondimensional trait value $w = \frac{r}{r_{\mathrm{max}}}$. Substituting in Equation S4 yields the following nondimensional model:

$$w(\tau) = \begin{cases} \left[ \left(\frac{\tau}{\alpha}\right)^{\alpha} \left(\frac{1-\tau}{1-\alpha}\right)^{1-\alpha} \right]^{s} & 0 < \tau < 1 \\ 0 & \text{otherwise} \end{cases} \tag{S6}$$

that removes all parameters that give units to the model. This shows that parameters $\alpha$ and $s$ determine the shape of the curve, while parameters $r_{\mathrm{max}}$,

$T_{\min}$ and $T_{\max}$ determine the location and scaling of the curve in the temperature and performance axes.

Currently, Equations S4, S5 and S6 have a parameter $s > 0$ that does not have a direct biological interpretation. However, differentiating Equation S6 yields

$$\begin{aligned}\frac{dw}{d\tau} &= w(\tau)\frac{d\ln w}{d\tau} \\ &= sw(\tau)\left[\frac{\alpha}{\tau} - \frac{1-\alpha}{1-\tau}\right]\end{aligned}$$

which shows that $s$ scales the magnitude of the rate of change of performance with temperature. Thus, large values of $s$ will give TPCs where performance decreases more steeply away from the optimum (i.e. curves that are more narrow).

To make the model more interpretable, it seems natural to try to find a way to substitute parameter $s$ by the thermal breadth of the TPC (i.e. the temperature range for which a certain relative performance reference value $w_{\text{ref}} \in (0, 1)$ is exceeded), a quantity of interest to ecologists. It is convenient to work with the nondimensional model to do this, as it only has two parameters and any derived results can easily be converted to analogous results for dimensional models. Mathematically, finding the (nondimensional) thermal breadth corresponds to finding $\tau_2 - \tau_1$, where $\tau_1, \tau_2$ are the solutions for the equation $w(\tau) = w_{\text{ref}}$, and $\tau_1 \le \tau_2$. Unfortunately, this equation does not have a closed-form solution. However, here we derive an approximate solution that is approached asymptotically as $w_{\text{ref}} \to 1$. To do this, we first take natural logarithms of Equation S6, substitute $w(\tau) = w_{\text{ref}}$ and divide by $s$ to get the equation

$$\frac{1}{s}\ln w_{\text{ref}} = \alpha \ln\frac{\tau}{\alpha} + (1-\alpha)\ln\frac{1-\tau}{1-\alpha}$$

with solutions $\tau_1, \tau_2$. Now, consider the second order Taylor expansion of the first term in the right hand side of this equation around $\tau = \alpha$. This yields

$$\alpha \ln\left(\frac{\tau}{\alpha}\right) \sim \tau - \alpha - \frac{1}{2\alpha}(\tau-\alpha)^2$$

Let $\rho = 1 - \tau$, $\gamma = 1 - \alpha$. After a change of variables and Taylor expanding in an analogous way, the second term can be approximated by

$$\begin{aligned}(1-\alpha)\ln\frac{1-\tau}{1-\alpha} = \gamma\ln\frac{\rho}{\gamma} &\sim \rho - \gamma - \frac{1}{2\gamma}(\rho-\gamma)^2 \\ &= \alpha - \tau - \frac{1}{2(1-\alpha)}(\alpha-\tau)^2\end{aligned}$$

Thus, as $\tau \to \alpha$ (which implies $w_{\text{ref}} \to 1$)

$$\begin{aligned}\frac{1}{s}\ln w_{\text{ref}} &\sim \tau - \alpha - \frac{1}{2\alpha}(\tau-\alpha)^2 + \left[\alpha - \tau - \frac{1}{2(1-\alpha)}(\alpha-\tau)^2\right] \\ &= -\frac{1}{2}\left(\frac{1}{\alpha} + \frac{1}{1-\alpha}\right)(\tau-\alpha)^2\end{aligned}$$

This is a quadratic equation on $\tau$, with solutions

$$\tau_1 = \alpha - \sqrt{-\frac{2\alpha(1-\alpha)}{s} \ln w_{\text{ref}}}$$
$$\tau_2 = \alpha + \sqrt{-\frac{2\alpha(1-\alpha)}{s} \ln w_{\text{ref}}}$$

and therefore, when $w_{\text{ref}} \to 1$ we have an asymptotic solution for the nondimensional thermal breadth $\tau_2 - \tau_1$:

$$\tau_2 - \tau_1 \sim \sqrt{-\frac{8\alpha(1-\alpha)}{s} \ln w_{\text{ref}}}$$

The choice of $w_{\text{ref}} = e^{-\frac{1}{8}}$ is particularly convenient, as in this case the expression simplifies to

$$\tau_2 - \tau_1 \sim \sqrt{\frac{\alpha(1-\alpha)}{s}}$$

We can define a new model parameter $\beta$ as an approximate nondimensional thermal breadth by replacing the asymptotic sign with equality so that

$$\beta = \sqrt{\frac{\alpha(1-\alpha)}{s}}$$

Solving for $s$, we get the expression

$$s = \frac{\alpha(1-\alpha)}{\beta^2}$$

which we can substitute into Equations S4 and S6 to get a fully interpretable parametrization of the flexTPC model in both nondimensional form

$$w(\tau) = \begin{cases} \left[ \left(\frac{\tau}{\alpha}\right)^{\alpha} \left(\frac{1-\tau}{1-\alpha}\right)^{1-\alpha} \right]^{\frac{\alpha(1-\alpha)}{\beta^2}} & 0 < \tau < 1 \\ 0 & \text{otherwise} \end{cases} \tag{S7}$$

and in what we consider the standard form of flexTPC

$$r(T) = \begin{cases} r_{\max} \left[ \left(\frac{T-T_{\min}}{\alpha}\right)^{\alpha} \left(\frac{T_{\max}-T}{1-\alpha}\right)^{1-\alpha} \left(\frac{1}{T_{\max}-T_{\min}}\right) \right]^{\frac{\alpha(1-\alpha)}{\beta^2}} & T_{\min} < T < T_{\max} \\ 0 & \text{otherwise} \end{cases} \tag{S8}$$

where the dimensional location/scaling parameters $T_{\min}, T_{\max}, r_{\max}$ are reintroduced, but the parameters $\alpha$, $\beta$ that determine the curve shape are kept in nondimensional form.

As before, we can also construct a fully dimensional parametrization in terms of $T_{\text{opt}}$ and the approximate thermal breadth $B$ at $r_{\text{ref}} = e^{-\frac{1}{8}} r_{\max}$ that may be of interest for applied scientists that wish to easily get estimates for these quantities from standard statistical software. The roots of $r(T) = r_{\text{ref}}$ can be obtained by transforming the dimensionless $\tau_1, \tau_2$ found previously into dimensional temperatures by inverting the equation $\tau = \frac{T - T_{\min}}{T_{\max} - T_{\min}}$, which leads to

$$\begin{aligned} T_1 &= T_{\min} + \tau_1 (T_{\max} - T_{\min}) \\ T_2 &= T_{\min} + \tau_2 (T_{\max} - T_{\min}) \end{aligned}$$

We can then get an asymptotic solution for the dimensional thermal breadth $T_2 - T_1$

$$\begin{aligned} T_2 - T_1 &= (\tau_2 - \tau_1)(T_{\max} - T_{\min}) \\ &\sim \beta (T_{\max} - T_{\min}) \end{aligned}$$

We can thus define $B = \beta(T_{\max} - T_{\min})$ as an approximate thermal breadth. Solving this expression for $\beta$ and substituting into Equation S7, along with the expressions that define the nondimensional quantities $w = \frac{r}{r_{\max}}$, $\alpha = \frac{T_{\text{opt}} - T_{\min}}{T_{\max} - T_{\min}}$, $1 - \alpha = \frac{T_{\max} - T_{\text{opt}}}{T_{\max} - T_{\min}}$, $\tau = \frac{T - T_{\min}}{T_{\max} - T_{\min}}$, $1 - \tau = \frac{T_{\max} - T}{T_{\max} - T_{\min}}$ yields the fully dimensional form of the flexTPC model

$$r(T) = \begin{cases} r_{\max} \left[ \left( \frac{T - T_{\min}}{T_{\text{opt}} - T_{\min}} \right)^{\frac{T_{\text{opt}} - T_{\min}}{T_{\max} - T_{\min}}} \left( \frac{T_{\max} - T}{T_{\max} - T_{\text{opt}}} \right)^{\frac{T_{\max} - T_{\text{opt}}}{T_{\max} - T_{\min}}} \right]^{\frac{(T_{\text{opt}} - T_{\min})(T_{\max} - T_{\text{opt}})}{B^2}} & T_{\min} < T < T_{\max} \\ 0 & \text{otherwise} \end{cases} \tag{S9}$$

The relative error for $B$ when approximating $T_2 - T_1$ is the same as the one for $\beta$ in approximating $\tau_2 - \tau_1$ (see Figure S4), since

$$\left| \frac{B - (T_2 - T_1)}{T_2 - T_1} \right| = \left| \frac{\beta(T_{\max} - T_{\min}) - (\tau_2 - \tau_1)(T_{\max} - T_{\min})}{(\tau_2 - \tau_1)(T_{\max} - T_{\min})} \right| = \left| \frac{\beta - (\tau_2 - \tau_1)}{\tau_2 - \tau_1} \right|.$$

### Constructing more parsimonious models from the flexTPC equation

FlexTPC is a five-parameter model that can describe thermal performance curves of a wide range of shapes. As all model parameters are interpretable, our preferred approach when data is limited is to take a Bayesian approach to parameter inference and use informative prior distributions based in biological knowledge (e.g. temperature ranges of the habitat of the species, or the approximate thermal breadth for previously fitted TPCs to the trait of interest in other species) to restrict parameter values to biologically reasonable ranges. This makes it possible to fit flexTPC even in data-limited situations. However, some researchers may not wish to take a Bayesian approach to parameter inference and may find a need for a more parsimonious model than flexTPC due to data limitations. In this section we show how the interpretable parametrization of flexTPC makes it possible to easily obtain models with fewer parameters from the flexTPC equation for these purposes.

To obtain a more parsimonious model, one or more of the flexTPC model parameters needs to be removed by either fixing it to a constant value or making it a function of the remaining model parameters. Whenever fitting a TPC to real data, it will almost always be necessary for the TPC model to be able to vary the maximum trait value $r_{\max}$ and the thermal limits $T_{\min}$ and $T_{\max}$ to describe the data. Because of this, the parameters that are the most likely candidates for removal are the parameter for the upper thermal breadth $\beta$ and the relative location of the thermal optimum $\alpha$ (although, as discussed in the main text, removing the latter will introduce a deterministic relationship between $T_{\text{opt}}$, $T_{\max}$, and $T_{\min}$ that can potentially introduce bias in estimates of these parameters).

##### Three parameter models

For very data-limited cases where introducing bias in the $T_{\text{opt}}$, $T_{\max}$, and $T_{\min}$ estimates is less important than having a parsimonious model that fits the data well, three-parameter TPC models may be useful. Various three parameter models can be obtained from the flexTPC equation by fixing both $\alpha$ and $\beta$.

For example, the quadratic model is a special case of flexTPC when fixing $\alpha = 1/2$, $\beta = 1/\sqrt{8}$. While the Briere1 model is not a special case of flexTPC, setting $\alpha \approx 0.8$, $\beta \approx 0.2$ yields a three-parameter model with a similar shape. Like the Briere1 model, this restricted flexTPC model also makes a strong assumption about the relationship of the temperature optimum and thermal limits. However, unlike the Briere1 model, this restricted model can describe TPCs from organisms that function at any temperature range, including below freezing temperatures.

##### Four parameter models

If estimating $T_{\text{opt}}$ and the thermal limits is one of the main goals of the analysis, researchers may wish to avoid possibly introducing bias into these quantities by introducing a deterministic relationship between these quantities, but still need a more parsimonious model than flexTPC. These kinds of models can be obtained by leaving $\alpha$ as a free parameter, and only removing $\beta$.

Rather than setting $\beta$ to a constant, a researcher may be interested in having the thermal breadth vary depending on curve skewness. We show that this can be done by making $\beta$ a function of $\alpha$. For example, suppose we want a model that reduces to the quadratic model for symmetric curves (i.e. $\beta = 1/\sqrt{8}$ when $\alpha = 0.5$) and approximates the Briere1 model for curves of similar skewness (i.e. $\beta = 0.2$ when $\alpha = 0.8$). A simple way of obtaining a restricted model from the flexTPC equation that accomplishes this is setting

$$\beta = \frac{1}{\sqrt{8}} - \frac{\left|\alpha - \frac{1}{2}\right|}{0.3}\left(\frac{1}{\sqrt{8}} - 0.2\right)$$

The resulting model has four free parameters ($T_{\text{min}}$, $T_{\text{max}}$,$r_{\text{max}}$ and $\alpha$) and can describe curves of any skewness (i.e. where $T_{\text{opt}}$ is at any point in-between $T_{\text{min}}$ and $T_{\text{max}}$), but assumes the thermal breadth varies deterministically with the skewness. This model can be used for traits for which either the Briere1 or quadratic models are typically used, making it possible to compare the inferred parameters fairly with a single four-parameter model.

### Shifted Briere model

In order to use the Briere model to describe TPCs for organisms that grow below freezing temperatures, the root at $T = 0°\text{C}$ can be shifted to a lower temperature by adding an offset temperature $T_{\text{offset}}$ to the function argument. In order to keep the interpretation of $T_{\text{min}}$ and $T_{\text{max}}$ as the minimum and maximum temperatures, the same offset needs to be added to these parameters. Modifying the Briere2 model equation

$$r(T) = cT(T - T_{\text{min}})(T_{\text{max}} - T)^{\frac{1}{m}}$$

by adding these offsets leads to the shifted Briere2 model

$$\begin{aligned} r(T) &= c(T + T_{\text{offset}})(T + T_{\text{offset}} - T_{\text{min}} - T_{\text{offset}})(T_{\text{max}} + T_{\text{offset}} - T - T_{\text{offset}})^{\frac{1}{m}} \\ &= c(T + T_{\text{offset}})(T - T_{\text{min}})(T_{\text{max}} - T)^{\frac{1}{m}} \end{aligned}$$

where now the root at $T = 0°\text{C}$ is replaced by a root at $T = -T_{\text{offset}}$. This model can describe TPCs for which $T_{\text{min}} \geq -T_{\text{offset}}$. However, the choice of $T_{\text{offset}}$ changes the shape of the resulting curve (see Figure S5). As before, a shifted Briere1 model corresponds to setting $m = 2$.

### Model comparison

We include alternative model comparison metrics to those used in the main text.

#### `botrana` dataset

##### Aikaike Information Criterion

| stage | Briere1 | Briere2 | flexTPC |
|---|---|---|---|
| eggs | **-55.89** | -40.25 | -49.61 |
| i1 | -59.19 | -54.92 | **-62.00** |
| i2 | -71.53 | -70.63 | **-74.22** |
| i3 | -41.38 | -33.46 | **-60.92** |
| i4 | -59.92 | **-63.21** | -58.29 |
| i5 | -57.53 | -75.16 | **-75.49** |
| pupae | -68.13 | -72.66 | **-98.48** |

**Bayesian Information Criterion**

| stage | Briere1 | Briere2 | flexTPC |
|---|---|---|---|
| eggs | **-53.33** | -37.06 | -45.78 |
| i1 | -56.64 | -51.73 | **-58.17** |
| i2 | -68.98 | -67.44 | **-70.39** |
| i3 | -38.82 | -30.26 | **-57.09** |
| i4 | -57.36 | **-60.02** | -54.45 |
| i5 | -54.97 | **-71.97** | -71.66 |
| pupae | -65.58 | -69.46 | **-94.65** |

#### `glacierbac` dataset

**Aikaike Information Criterion**

| species (strain) | Briere1 | Briere2 | flexTPC |
|---|---|---|---|
| *Arthrobacter sp* (55) | -60.66 | -63.68 | **-67.31** |
| *Arthrobacter sp* (60) | -37.43 | -46.32 | **-56.29** |
| *Pseudomonas* (76) | -31.86 | -30.17 | **-36.84** |
| *Arthrobacter glacialis* (137) | -41.60 | -38.82 | **-50.41** |
| *Arthrobacter glacialis* (158) | -61.54 | -66.20 | **-90.24** |

**Bayesian Information Criterion**

| species (strain) | Briere1 | Briere2 | flexTPC |
|---|---|---|---|
| *Arthrobacter sp* (55) | -59.07 | -61.69 | **-64.92** |
| *Arthrobacter sp* (60) | -36.64 | -45.33 | **-55.10** |
| *Pseudomonas* (76) | -31.07 | -29.19 | **-35.65** |
| *Arthrobacter glacialis* (137) | -41.82 | -39.09 | **-50.74** |
| *Arthrobacter glacialis* (158) | -60.75 | -65.21 | **-89.06** |

##### `abcoli` dataset

###### Aikaike Information Criterion

| antibiotic | Briere1 | Briere2 | flexTPC |
|---|---|---|---|
| AMP | -5.56 | -8.64 | **-17.12** |
| CLI | **-11.98** | -10.51 | -11.38 |
| CPR | -2.92 | -7.16 | **-11.75** |
| ERY | 3.28 | 0.25 | **-4.95** |
| FOX | -18.14 | -16.31 | **-21.11** |
| GEN | -3.16 | -3.96 | **-11.56** |
| LVX | **-12.93** | -10.96 | -12.43 |
| NTR | -16.03 | -16.47 | **-17.42** |
| STR | -5.39 | -7.42 | **-12.87** |
| TET | **-15.07** | -13.83 | -14.88 |
| TMP | -18.63 | -16.74 | **-24.29** |
| TOB | -7.15 | -9.40 | **-17.35** |
| no drug | -5.49 | -6.43 | **-25.33** |

###### Bayesian Information Criterion

| antibiotic | Briere1 | Briere2 | flexTPC |
|---|---|---|---|
| AMP | -5.78 | -8.91 | **-17.45** |
| CLI | **-12.19** | -10.78 | -11.70 |
| CPR | -3.14 | -7.43 | **-12.07** |
| ERY | 3.07 | -0.02 | **-5.27** |
| FOX | -18.35 | -16.58 | **-21.43** |
| GEN | -3.38 | -4.23 | **-11.88** |
| LVX | **-13.15** | -11.23 | -12.75 |
| NTR | -16.25 | -16.74 | **-17.74** |
| STR | -5.61 | -7.69 | **-13.19** |
| TET | **-15.28** | -14.10 | -15.20 |
| TMP | -18.85 | -17.01 | **-24.61** |
| TOB | -7.37 | -9.67 | **-17.68** |
| WT | -5.70 | -6.70 | **-25.65** |

#### Statistical model for Bayesian estimation of temperature-dependent mosquito life history traits

For each trait in the `lhculex` dataset, we compare the flexTPC model to a different TPC model that was used to fit the same data in a previous study (Shocket et al, 2020).

| Species | Egg viability | Larval survival | Development rate | Adult lifespan |
|---|---|---|---|---|
| *Culex pipiens* | Quadratic | Quadratic | Briere1 | Linear |
| *Culex quinquefasciatus* | Briere1 | Quadratic | Briere1 | Linear |

Table 1: Previously used model in the literature for each trait and species.

where the models are defined as follows:

**Briere1:** $r(T) = \begin{cases} cT(T - T_{\min})(T_{\max} - T)^{\frac{1}{2}} & T_{\min} < T < T_{\max} \\ 0 & \text{otherwise} \end{cases}$

**Quadratic:** $r(T) = \begin{cases} c(T - T_{\min})(T_{\max} - T) & T_{\min} < T < T_{\max} \\ 0 & \text{otherwise} \end{cases}$

**Linear:** $r(T) = \begin{cases} m(T_{\max} - T) & T > 15°\text{C} \\ m(T_{\max} - 15) & \text{otherwise} \end{cases}$

(note that the "linear" model for lifespan is really a piecewise linear model where lifespan is assumed constant below 15°C).

##### Model reparametrization

In order to make comparisons between the models fair, identical prior distributions were used for the parameters of both flexTPC and the corresponding literature model whenever possible. This required reparametrizing the quadratic, Briere1 and linear models so they can be written in terms of the maximum trait value $r_{\max}$.

The "linear model" that was used in Shocket et al for adult mosquito lifespan is in truth a piecewise linear model that is assumed to be constant below 15°C and is linear above this temperature. Since lifespan decreases at high temperatures, this means that in this model $r_{\max} = m(T_{\max} - 15)$. We can use this expression to reparametrize the linear model and write it down in terms of $T_{\max}$ and $r_{\max}$:

$$r(T) = \begin{cases} r_{\max}\left(\frac{T_{\max} - T}{T_{\max} - 15}\right) & T > 15°\text{C} \\ r_{\max} & \text{otherwise} \end{cases}$$

The quadratic model is a special case of the flexTPC model if we set $\alpha = \frac{1}{2}$ and $s = 2$ in Equation S4. This yields the following expression:

$$r(T) = \begin{cases} \frac{4r_{\max}}{(T_{\max} - T_{\min})^2}\left(T - T_{\min}\right)\left(T_{\max} - T\right) & T_{\min} < T < T_{\max} \\ 0 & \text{otherwise} \end{cases}$$

which is written down in terms of $r_{\max}$.

The Briere1 model cannot be written down in terms of $r_{\max} = r(T_{\text{opt}})$ in a simple way, but if the parameters $r_{\max}$, $T_{\min}$ and $T_{\max}$ are known, we can solve the Briere1 model equation for $c$ to get

$$c = \frac{r_{\max}}{T_{\text{opt}}(T_{\text{opt}} - T_{\min})(T_{\max} - T_{\text{opt}})^{\frac{1}{2}}}$$

where $T_{\text{opt}} = \frac{4T_{\max}+3T_{\min}+\sqrt{16T_{\max}^2+9T_{\min}^2-16T_{\min}T_{\max}}}{10}$. We can thus set a prior distribution on parameters $r_{\max}, T_{\min}, T_{\max}$ for the Briere model and calculate $c$ as a transformed parameter.

In the following sections, we use $\mathcal{P}$ as a notational shortcut to denote all parameters of the correponding TPC model. For example, for the Briere1 and quadratic models we have $\mathcal{P} = \{T_{\min}, T_{\max}, r_{\max}\}$ while for the flexTPC model we have $\mathcal{P} = \{T_{\min}, T_{\max}, r_{\max}, \alpha, \tilde{\beta}\}$. We use the flexTPC model as parametrized in Equation S8 for inference.

#### Probability of larval survival to adulthood

As no information was easily available about the number of mosquitoes used to calculate the observed proportions in the dataset, we used a normal likelihood rather than a binomial for this dataset.

##### Likelihood

$$p_i | T_i, \mathcal{P} \sim \text{Normal}(\mu = r(T_i : \mathcal{P}), \sigma)$$

##### Prior distributions

The following priors were used for the flexTPC model:

$$\begin{aligned}
T_{\min} &\sim \text{Normal}(\mu = 5, \sigma = 2.5) \\
T_{\max} &\sim \text{Normal}(\mu = 35, \sigma = 5) \\
r_{\max} &\sim \text{Uniform}(0, 1) \\
\alpha &\sim \text{Uniform}(0, 1) \\
\beta &\sim \text{Gamma}(\mu = 0.2, \sigma = 0.4) \\
\sigma &\sim \text{Uniform}(0, 1)
\end{aligned}$$

The same priors for $T_{\min}, T_{\max}$, $r_{\max}$ and $\sigma$ were used for the quadratic model.

The same priors for $T_{\min}, T_{\max}$ and $r_{\max}$ were used for the Briere1 (for *Culex pipiens*) and quadratic (for *Culex quinquefasciatus*) models. The priors for the minimum and maximum temperatures are weakly informative, and correspond to *a priori* assumptions that the minimum temperature is $\approx 95\%$ likely to be in the temperature range $[0°\text{C}, 10°\text{C}]$, that the maximum temperature is $\approx 95\%$ likely to be in the temperature range $[30°\text{C}, 40°\text{C}]$, which are reasonable for the mosquito species. The prior for the maximum trait value is noninformative, assuming that the maximum proportion of viable eggs is *a priori* equally likely to be in the interval $[0, 1]$. The prior for the thermal breadth parameter $\beta$ is (very) weakly informative. It has a mean of 0.2 (corresponding to a similar thermal breadth as the Briere1 model) and a 95% prior credible interval of $[2.1 \times 10^{-7}, 1.37]$. It places higher prior probability on temperature breadths that are similar to those in commonly used TPC models, like the Briere1 and quadratic models (for reference, for the quadratic model $\beta = \frac{1}{\sqrt{8}} \approx 0.35$), but allows for other values if necessary to describe the data. The prior for the standard deviation is uniform, with an upper bound that is higher than values that could be taken by the data (which is bounded to be in the interval $[0, 1]$).

#### Mosquito development rate

##### Likelihood

$$y_i | T_i, \mathcal{P} \sim \text{Normal}(\mu = r(T_i; \mathcal{P}), \sigma)$$

##### Prior distributions

The following priors were used for the flexTPC model:

$$\begin{aligned}
T_{\text{min}} &\sim \text{Normal}(\mu = 5, \sigma = 2.5) \\
T_{\text{max}} &\sim \text{Normal}(\mu = 35, \sigma = 5) \\
r_{\text{max}} &\sim \text{Uniform}(0, 1) \\
\alpha &\sim \text{Uniform}(0, 1) \\
\beta &\sim \text{Gamma}(\mu = 0.2, \sigma = 0.4) \\
\sigma &\sim \text{Uniform}(0, 1)
\end{aligned}$$

The same priors for $T_{\text{min}}, T_{\text{max}}$, $r_{\text{max}}$ and $\sigma$ were used for the Briere1 model.

The prior distribution for $r_{\text{max}}$ needs a special note, as the response is not a probability and as such is not bounded at one mathematically. The prior for $r_{\text{max}}$ now corresponds to an assumption that the mosquito development rate (i.e. the inverse of the time to adulthood in days) is *a priori* equally likely to be in the interval $[0, 1]$. The upper bound is appropriate because the time from egg hatching to adulthood will be substantially more than a day (resulting on MDR values less than one) even under optimal conditions. The reasoning for the priors used for the other model parameters is similar as for those mentioned previously for the probability of larval survival.

#### Adult lifespan

##### Likelihood

$$y_i | T_i, \mathcal{P} \sim \text{Normal}(\mu = r(T_i; \mathcal{P}), \sigma)$$

##### Prior distributions

The following priors were used for the flexTPC model:

$$\begin{aligned}
T_{\min} &\sim \text{Normal}(\mu = 5, \sigma = 2.5)\\
T_{\max} &\sim \text{Normal}(\mu = 35, \sigma = 5)\\
r_{\max} &\sim \text{Uniform}(0, 150)\\
\alpha &\sim \text{Uniform}(0, 1)\\
\beta &\sim \text{Gamma}(\mu = 0.2, \sigma = 0.4)\\
\sigma &\sim \text{Uniform}(0, 100)
\end{aligned}$$

The same priors for $T_{\max}$, $r_{\max}$ and $\sigma$ were used for the linear model.

The reasoning for the prior distributions for $T_{\min}$, $T_{\max}$, $\alpha$ and $\beta$ are the same as before. The prior for $r_{\max}$ corresponds to the assumption that the maximum adult lifespan (in days) is a priori equally likely to be in the interval $[0, 150]$. Due to the increased range of the response variable, the upper bound for the uniform prior for the standard devation was also increased.

#### Egg viability

As information on the number of eggs used to calculate the proportions at each temperature was available for this dataset, egg viability was modeled with a binomial distribution for the number of viable eggs $n_i$ out of $N_i$ total eggs that were evaluated for viability at temperature $T_i$.

##### Likelihood

$$n_i | T_i, \mathcal{P} \sim \text{Bin}(p = r(T_i; \mathcal{P}), N_i)$$

##### Prior distributions

The following priors were used for the flexTPC model:

$$\begin{aligned}
T_{\min} &\sim \text{Normal}(\mu = 10, \sigma = 5)\\
T_{\max} &\sim \text{Normal}(\mu = 35, \sigma = 5)\\
r_{\max} &\sim \text{Uniform}(0, 1)\\
\alpha &\sim \text{Uniform}(0, 1)\\
\beta &\sim \text{Gamma}(\mu = 0.2, \sigma = 0.4)
\end{aligned}$$

The same priors for $T_{\min}, T_{\max}$ and $r_{\max}$ were used for the Briere1 (for *Culex pipiens*) and quadratic (for *Culex quinquefasciatus*) models. The prior distribution for $T_{\min}$ was made less informative than in the other traits, as initial attempts to fit this trait showed that the $T_{\min} \sim \text{Normal}(\mu = 5, \sigma = 2.5)$ prior used for other traits was too restrictive, as $T_{\text{min}}$ values above 10°C are reasonable for this trait. Thus, the prior distribution was modified so that the prior 95% credible interval was approximately $[0°\text{C}, 20°\text{C}]$. The reasoning for the prior distributions of other parameters was identical as detailed above for other traits.

### References

[1] Mauricio Cruz-Loya, Elif Tekin, Tina Manzhu Kang, Natalya Cardona, Natalie Lozano-Huntelman, Alejandra Rodriguez-Verdugo, Van M. Savage, and Pamela J. Yeh. Antibiotics Shift the Temperature Response Curve of Escherichia coli Growth. *mSystems*, page e0022821, July 2021.

[2] Xinyou Yin, Martin J. Kropff, Graham McLaren, and Romeo M. Visperas. A nonlinear model for crop development as a function of temperature. *Agricultural and Forest Meteorology*, 77(1):1–16, November 1995.
